# Automated measurement of cardiomyocyte monolayer contraction using the Exeter Multiscope

**DOI:** 10.1101/2024.09.09.611998

**Authors:** Sharika Mohanan, David Horsell, Taylor Watters, Mohammadreza Ghasemi, Lewis Henderson, Caroline Müllenbroich, Gil Bub, Francis Burton, Godfrey Smith, Alexander D. Corbett

## Abstract

We apply a novel microscope architecture, the Exeter Multiscope, to the problem of acquiring image data in rapid succession from nine wells of a 96 well plate. We demonstrate that the new microscope can detect contraction in cardiomyocyte monolayers which have been plated into these wells. Each well is sampled using 500 × 500 pixels across a 1.4 × 1.4 mm field of view, acquired in three colours at 3.7 Hz per well. The use of multiple illumination wavelengths provides post-hoc focus selection, further increasing the level of automation. The performance of the Exeter Multiscope is benchmarked against industry standard methods using a commercial microscope with a motorised stage and demonstrates that the Multiscope can acquire data almost 40 times faster. The data from both Multiscope and the commercial systems are processed by a ‘pixel variance’ algorithm that uses information from the pixel value variability over time to determine the timing and amplitude of tissue contraction. This algorithm is also benchmarked against an existing algorithm that employs an absolute difference measure of tissue contraction.

## INTRODUCTION

Within industrialised nations, one in two people in will die of heart disease. Our ability to screen for new drugs to prevent more cases and improve outcomes is severely limited by the rate at which assays can be performed. There exists a bottleneck in testing candidate drugs using standard biological models such as cardiomyocyte monolayers. Part of the bottleneck problem is in obtaining imaging data to assess drug performance at various concentrations in cardiomyocyte monolayers over many days and weeks. These assay experiments often take place in the wells of a standardised multiwell plate. However, to image the wells, they must first be individually translated into the imaging path of an optical microscope. For 96-and 384-well plates, this translation can be very time consuming, even with the use of motorised stages and automated control software.

Previous work to address the problem of throughput in high content imaging has focused on imaging the entire well plate area onto the camera detector^1,2^. This greatly improves temporal resolution, but at the cost of spatial resolution per well (i.e. sampling at >100 um per pixel). There have also been the challenges in obtaining uniform illumination across the well plate and maintaining telecentric imaging across such large areas.

The Exeter Multiscope uses a very different approach to parallelise image acquisition across each well in a multiwell plate. Based on the concept first published as the Random Access Parallel (RAP) microscope^3^, the Exeter Multiscope (MS) constructs a complete imaging system around each well in a 96-well plate. The 9 mm pitch between wells limits the size and complexity of the illumination and detection optics that can be employed, but we demonstrate that these compromises do not limit our ability to detect tissue contraction. In this study we demonstrate that it is possible to detect tissue contraction in nine wells in rapid succession, with the potential to extend this to many more. The MS contraction data was benchmarked against data acquired within Clyde Biosciences Ltd. (CB), a contract research organisation that provides an assay service for pharmaceutical companies to provide clinically relevant pharmacological data using cardiomyocyte monolayer models.

In addition to identifying differences in data acquisition, two separate image processing algorithms were used. The first, MuscleMotion (MM) identifies contraction peaks by calculating the difference in pixel values at two different time points and taking the modulus of these values. A second method, described as the Pixel Variance (PV), method calculates the standard deviation in pixel values for all frames over a defined time interval. Both the PV and MM algorithms are applied to both the data acquired by the Clyde Biosciences microscope and the Exeter Multiscope.

## METHODS

### The Exeter Multiscope (MS) prototype

A sketch showing the optical layout of the Multiscope is shown in Figure 1(A). This is based on the original Random Access Parallel (RAP) microscope design first described in reference 3. The central feature of the design is that any light that passes through the wells is directed to the same point (the exit pupil of the system) where an area detector is located. This is achieved using a final steering optic in the light path. Each well in the well plate is imaged using its own individual light emitting diode (LED) light source, collimation lens and imaging objective, all of which must all have a lateral extent smaller than the 9 mm well pitch. Switching on one LED will relay an image of the sample above it to the camera. Unlike alternative single camera high content imaging systems, which image the entire well plate onto the camera, the image of each well is sampled by the full resolution of the camera chip. This trades temporal resolution for spatial resolution. The trade-off can be tapered by selecting the field of view of each well image, with smaller fields of view enabling higher camera frame rates.

**Figure 1.**
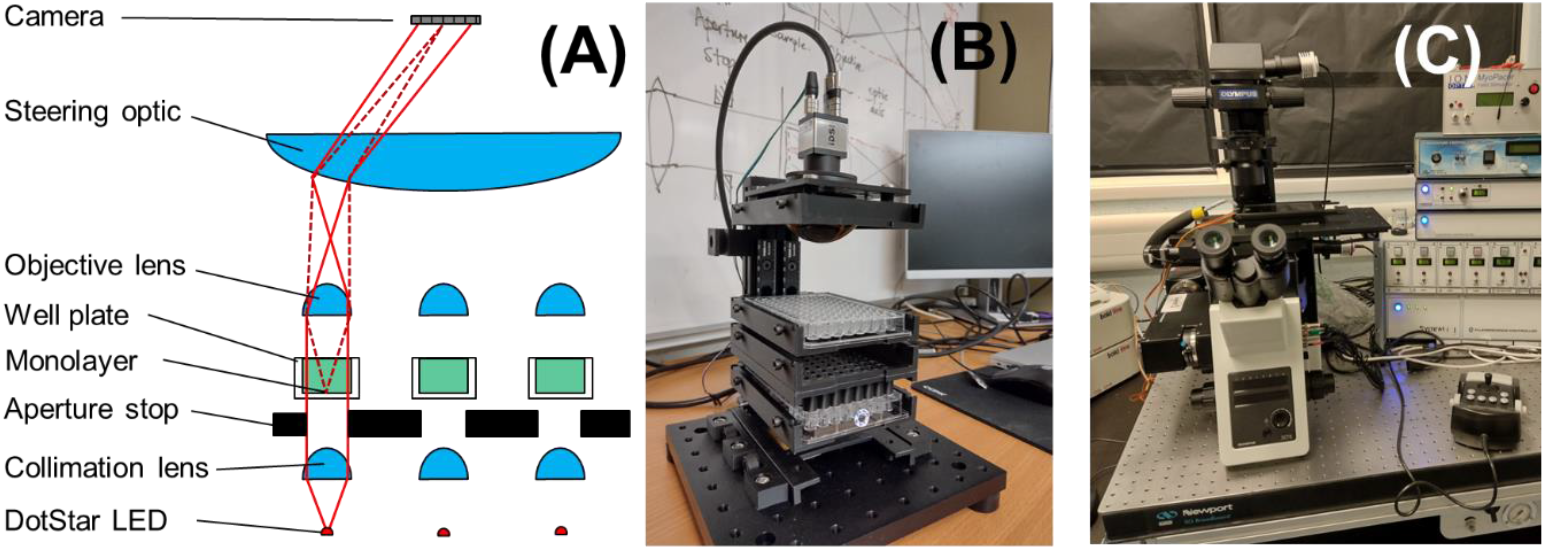
(A) Outline of the Multiscope optical design indicating how multiple fields of view can be relayed to the same exit pupil. (B) Image of the Multiscope prototype (C) Image of the Clyde Biosciences motorised microscope.

A second central feature of the Multiscope is that the illumination is collimated. Light from the small LED emitter area (<300 µm × 300 µm), when collimated by an array of 6 mm Ø lenses, produces a beam divergence of less than 0.3 mrad. Illumination by strongly collimated light provides a form of phase contrast imaging (Schlieren imaging) that converts phase gradients to intensity values. This is because light leaving the sample at angles higher than 5.7° do not pass the small angular aperture of the objective lenses and are filtered out of the image. The high contrast of the Schlieren images ensures that the distortion and warping from the monolayer contraction can be detected with high sensitivity. The fact that the sample illumination is collimated rather than convergent results in the separation of the illumination path (shown by solid lines in Fig 1(A)) and the imaging path (dotted line, Figure 1(A)).

The first key challenge in constructing the Multiscope is obtaining LED arrays that are individually addressable and can be located on the same 9 mm pitch as the wells in a 96-well plate. In this prototype, standalone DotStar LEDs were used (SK9822, Adafruit) which are contained in a 5 mm x 5 mm package. Mechanical alignment of a 3 × 3 array of LEDs was achieved by 3D printing a plate into which recesses had been made which were just large enough to house the LED package. The LED was pushed into the recess emitter-side first with small apertures in the recesses allowing the light to pass through the plate. The underside of the LEDs were then fully accessible to allow data and power connections to be wired and hand soldered.

The alignment of the collimation lenses was achieved using an empty, flat-bottomed well plate (Greiner 655101) into which Thorlabs uncoated, 30 mm focal length plan-convex lenses (LA1700) could be placed flat-side down. As the lenses diameter is only fractionally smaller than the well diameter (6.0 mm vs 6.39 mm) no further lateral adjustment of the lenses was required. The same array of nine 30 mm focal length collimation lenses was repeated to form the array of objective lenses above the sample well plate.

The light from each of the DotStar LEDs has a very broad angular distribution. It is then possible for stray light from one LED to reach the collimation lens of an adjacent well position. After passing through the optical path of the adjacent well, this stray light could then produce a dim image of the adjacent sample on top of the brighter image of the intended sample. To avoid this cross-talk, a series of 10 mm deep aperture stops were introduced between the collimation lens and the sample. These aperture stops were produced from black plastic multiwell plates (Scientific Laboratory Supplies Limited, Sterilin Microplate Black 611F96BK) in which the base had been drilled out, although custom aperture stops could equally be 3D printed.

To maintain the lateral and axial alignment of the first five layers of optics, plate holders were 3D printed which supported the well plates on three sides. All plate holders were attached to the same pair of dovetail rails (Thorlabs RLA300/M) via two rail sliders (Thorlabs RC1) to provide vertical adjustment. Small lateral adjustments of the plates could be made through the use of two opposing pairs of M3 screws introduced into the long sides of the plate holders via threaded holes.

The large steering optic was provided by a 75 mm diameter, 60 mm focal length condenser lens (Thorlabs ACL7560U-A). Whilst not designed for imaging, the purpose of the condenser lens is to steer the light from each objective lens towards a common exit pupil for the system, which is coplanar with the camera chip. Whilst off-axis illumination of the condenser lens does introduce some image distortion, in this application it is only changes between images (through tissue contraction) that are of interest. The condenser lens was placed in a mount (Thorlabs LMR75) and the mount screwed into a 3D printed adaptor plate, which had the same dimensions as a well plate. This adaptor plate then sat within the same plate holder used to align the other well plates. Finally, the camera was attached to a Thorlabs mount (SM2A55, SM2F1) which was then screwed into a 3D printed plate which was attached directly to the condenser lens adaptor plate to maintain alignment between the camera and the condenser lens.

Each of the DotStar LEDs is a cluster LED, providing red, green and blue (R, G, B) emitters (SK9822, Adafruit). These emitters are closely spaced, ensuring the fields of illumination for each colour overlap completely across the entire field of view. The LED colour can be individually addressed and switched in less than 10 ns. The simple plano-convex lenses used to image each well have no chromatic correction. The inherent longitudinal chromatic aberration in these lenses means that each illumination colour focuses at a different depth within the sample. This ‘chromatic focusing’ effect was exploited to circumvent the need to manually focus each sample. According to simulation (ZEMAX OpticsStudio, Ansys Inc.), the focal planes of the LA1700 lens are shifted relative to the blue channel by 0.21 mm (green) and 0.46 mm (blue). Together with the simulated 0.22 mm depth of field of the condenser lens, the total depth range covered by chromatic focusing is expected to be 0.68 mm. This value is expected to exceed the anticipated separation between the focal plane and sample due to the mechanical tolerances of the MS tray position, the 1% tolerance in the focal length of the objective lenses, and variations in the precise dimensions of the well plate. During data collection, images of each well were captured under illumination by each colour before moving on to the next well. Capturing images of each well using red, green and blue illumination meant the data could be analysed post-hoc to find the illumination wavelength which focused closest to the monolayer sample.

Control of the illumination sequence was coordinated by a Raspberry Pi Pico (Pimoroni). The Pico used a GPIO connection to the machine vision camera (IDS UI-3060CP-M-GL Rev.2) to detect output TTL pulses that signalled the triggering of a new frame. Once the trigger was detected, the Pico addressed the LEDs to illuminate the next well, with the first colour in the (R, G, B) sequence. After illuminating the well with all three colours in three separate frames, the next well was illuminated in the same sequence to capture three more frames. This continued until all nine wells had been captured in all three colours, after which the sequence started again.

### The Clyde Biosciences (CB) microscope

The Clyde Biosciences microscope is constructed around a standard Olympus IX73 inverted microscope. The samples were illuminated by a white LED (Cairn OptoLED) with the brightfield images captured by an sCMOS camera (Hamamatsu ORCA Flash 4.0). Lateral and axial locations of the best focus images for each of the nine wells were manually obtained by using a motorised stage (Prior Scientific UK) to visit each well position and record each of the lateral and axial positions into the automated control software. Image data was acquired using a 40X 0.8 NA lens from each well for 700 frames at a sampling rate of 100 Hz before moving to the next well position. A comparison table of the CB and MS systems is provided in Table 1.

**Table 1:** Comparison of the Multiscope (MS) and Clyde Biosciences (CB) microscope properties.

| Component | Parameter | Clyde Biosciences (CB) | Multiscope (MS) |
| --- | --- | --- | --- |
| Sample environment |  |  |  |
|  | Temperature control | Incubator | None |
|  | CO2 control | Incubator | None |
| Imaging |  |  |  |
| | Source | LED (Incoherent) | μLED (Semi coherent) |
| | Wavelengths | Broadband | 465, 520, 625 ± 5 nm |
|  | Imaging modality | Brightfield | Phase contrast |
|  | Magnification | 40X | 2X |
|  | Numerical aperture | 0.8 | 0.1 |
|  | Theoretical resolution XY | 386 nm | 3000 nm |
| | Field of view (after crop) | 82 x 72 μm | 1465 x 1465 μm |
|  | Sample focusing | Manual | Chromatic |
|  | ROI selection | Manual | Fixed |
|  | Stage | Motorised | Fixed |
|  | # accessible wells | 96 | 9 |
| Detection |  |  |  |
|  | Frame rate per well | 100 fps / well | 100, 11, 3.7 fps / well |
| | Camera pixel size | 6.5 μm | 5.86 μm |
| | Spatial sampling | 0.16 μm/pixel | 2.9 μm/pixel |
|  | Sampling / Nyquist | 1.2 x Nyquist | 0.5 x Nyquist |
|  | Image size | 512 x 450 pixels | 500 x 500 pixels |
|  | Detector area used | 3.3 x 2.9 mm | 11.3 x 7.12 mm |
| Data processing |  |  |  |
|  | Analysis software 1 | MuscleMotion (MM) | MuscleMotion (MM) |
|  | Analysis software 2 | Pixel variance (PV) | Pixel variance (PV) |

A key difference between the Exeter Multiscope (MS) and Clyde Biosciences (CB) systems is the integration of a stage incubator on the CB system. The stage incubator controls the temperature, humidity and CO_2_ content of the monolayer environment, minimising variability in contraction behaviour between experiments. By contrast, the MS system does not have any environmental control, meaning that the monolayer tissue experiences rapid cooling after removal from the 37 °C stage incubator to the 21°C room temperature before MS imaging. This rapid cooling is known to significantly reduce the spontaneous beat rate of the tissue. The rapid reduction in temperature before MS imaging also introduced condensation for several seconds after transfer, which acts to diffuse the collimated illumination. Only after the condensation has coalesced can the imaging begin.

The imaging objective of the CB system has an NA of 0.8, compared to the 0.1 NA of the MS (Figure 2). In addition to a much higher spatial resolution, this provides a much smaller depth of field, reducing the influence of small bubbles or circulating particles outside of the focal plane. By contrast the MS system has an NA of 0.1, which had a much larger depth of field and included much of this out-of-plane movement, which interfered with the detection of monolayer contraction events. The low NA of the MS enabled much larger field of view imaging, making it less susceptible to data lost due to local gaps in the monolayer or regions of poor adhesion to the well plate. By capturing data from multiple focal planes using the chromatic focusing effect, the data from the MS could also be acquired automatically, without human intervention or mechanical movement of the tissue or optics.

**Figure 2:**
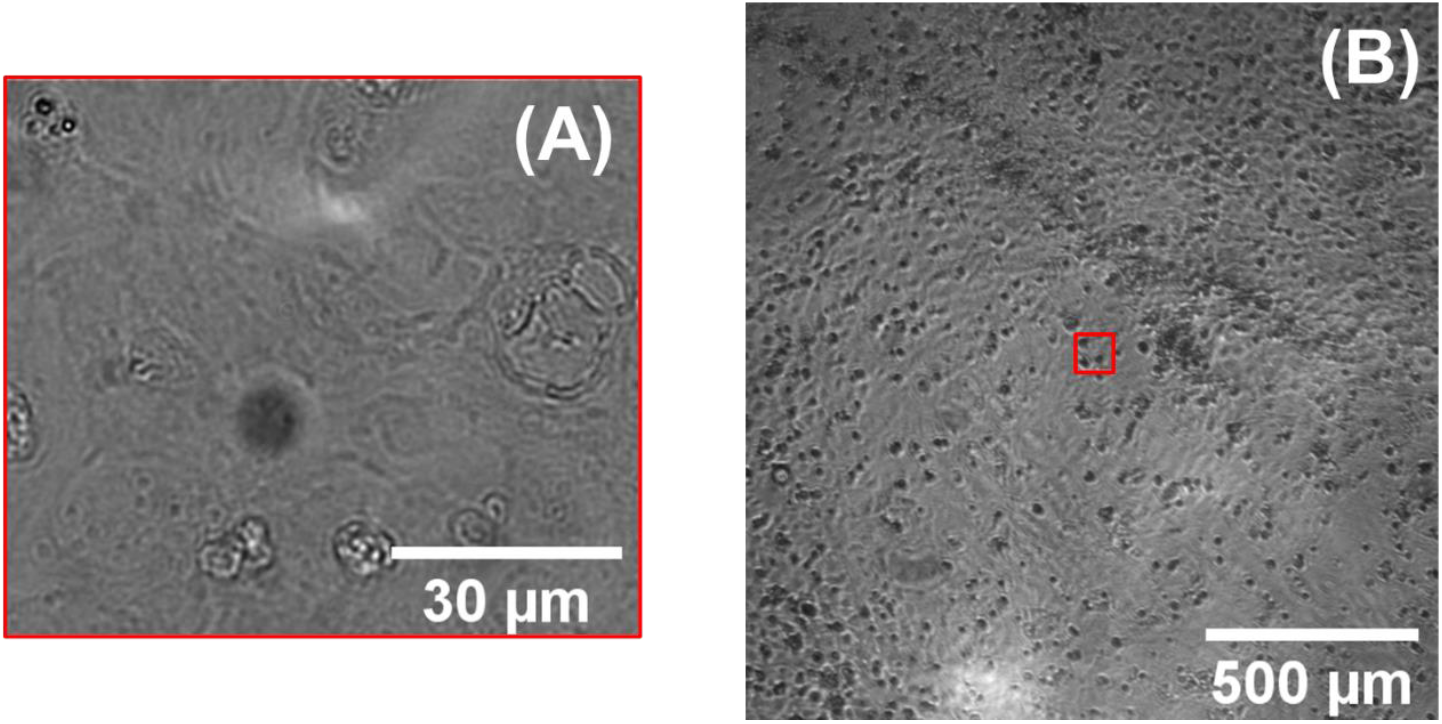
Example cardiomyocyte monolayer images acquired with Clyde Biosciences (A) and Multiscope (B) microscopes. The size of the field of view of the CB microscope is represented by a red square in (B).

### Human-Induced Pluripotent Stem Cell–Derived Cardiomyocyte Cell Culture iCell^**2**^

Cardiomyocytes (FUJIFILM Cellular Dynamics, Madison, WI, USA) were kept at −190 °C and prepared for culture as per the manufacturer’s instructions. Cell donor (01434) was registered with the ethics committee for research uses (NICHD-NIH, USA, with approval number N-01-HD-4-2865). Standard fibronectin vs. Matrix Plus well coating: The cells were cultured using two different coatings in a humidified incubator at 37 °C with 5% CO_2_. First, 96-well glass-bottomed plates (MatTek, Ashland, MA) were coated with fibronectin (10 mg/mL in PBS supplemented with Ca^2+^ and Mg^2+^; Sigma, St. Louis, MO, USA) (defining FM). Second, CELLvo™ Matrix Plus (StemBioSys, San Antonio, TX, USA), which were pre-coated on 96 well glass bottom plates and kept at 4 °C. On the day of plating, CELLvo™ Matrix Plus plates were transferred to room temperature, rehydrated with phosphate buffer saline (PBS), incubated for 1 h at 37 °C, and washed 2× with PBS. Cells were introduced at a density of 50,000 cells/well to allow the formation of a confluent monolayer in each well. The maintenance protocols followed the manufacturer’s instructions and used the iCell^2^ Cardiomyocytes Maintenance media for media changes every 2 days. Experiments were performed between days 5 and 6, as recommended by the manufacturers. Before beginning an experiment, cells were washed in serum-free media (SF media) (Fluorobrite DMEM, Gibco, Thermo Fisher Scientific, Horsham, UK).

### Data capture and processing

After the raw images have been acquired, contraction data was extracted using two different algorithms. The first, MuscleMotion (MM) is based on an absolute difference measure of image change^4^. This algorithms builds on previous methods to image tissue contraction^5–7^. In brief, a pixel-wise subtraction is performed between images acquired at two different time points, separated by a known number of frames (the ‘frame delay’). The modulus value of this image subtraction is calculated and then the pixel values are summed across the entire processed image. In this way, a large change in pixel value, experienced by most image pixels, between time points will yield the largest change in the absolute difference metric.

The modulus difference can be calculated as a running subtraction i.e. with a fixed frame delay between the current frame and the previous comparison frame (Figure 3). This provides the ‘contraction speed’ and provides a measure of the rate of contraction. As the sign of the image difference is lost by the modulus step, the direction of the image change is also lost. This means that the absolute displacement of the contraction cannot be determined by integration of the contraction speed metric. To obtain the absolute contraction displacement, the absolute difference metric must be calculated relative to a fixed reference point in the time sequence. This could be the first image frame or an image frame in which the tissue is known to be quiescent. When comparing the ‘contraction’ and ‘contraction speed’ profiles calculated in this way, the latter should appear (at least qualitatively) like the time derivative of the first. The MM software is free to use and can be implemented as a java plugin for ImageJ/Fiji.

**Figure 3:**
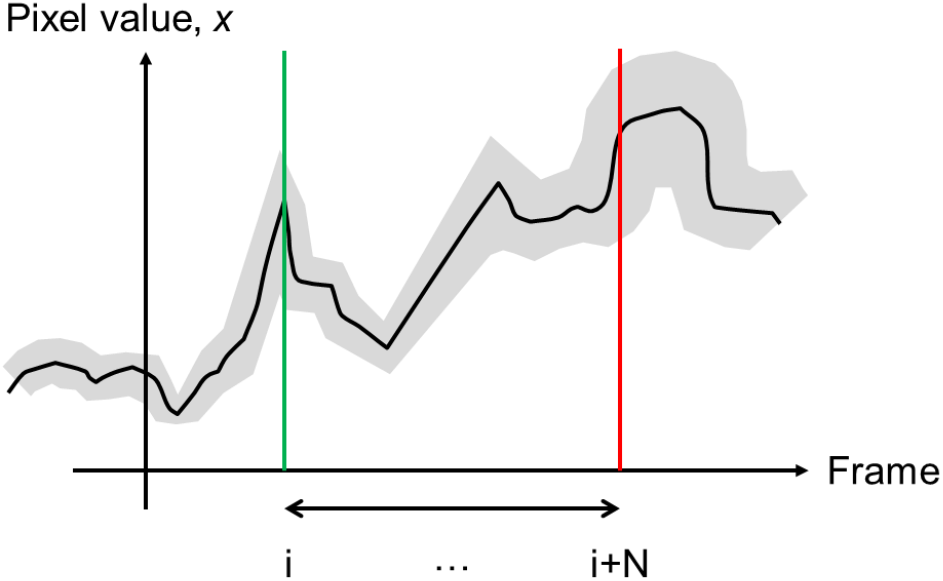
Sketch showing how the value of a single monochrome pixel may appear over time (black line). The Poisson statistics of the detection introduce a variance in the measured pixel value which scales linearly with the pixel value (indicated by the grey region). MM analysis calculates the modulus of the difference in the pixel value between two time points separated by the frame delay (N frames). PV analysis calculates the variance of the pixel values at all time points within the frame delay interval.

The MM software has many additional features which enable the extraction of numerous parameters associated with contraction (rise time, contraction duration etc.) but these will not be discussed in detail here. There are essentially two outputs of the MM software: the ‘contraction speed’ and the ‘absolute contraction’.

### Pixel Variance (PV)

The Pixel Variance (PV) method of image processing acts in a similar way to the MM algorithm and has been applied successfully in the past to cardiac data^8^. The key difference is that when calculating a ‘contraction speed’ measurement, the algorithm uses not only the data from the two frames separated by the known frame delay, but from all of the frames in between. For each pixel location in the image, the pixel value at each time point over the frame delay interval is extracted. The standard deviation of the pixel values is then calculated and becomes the value of the pixel in the processed image. By using all the time points within the frame interval rather than just two, it is expected that any noise in the metric will reduce as the frame delay increases, rather than remain fixed as it does for the modulus subtraction method (see Appendix B). A visual summary of the process is given in Figure 4.

**Figure 4:**
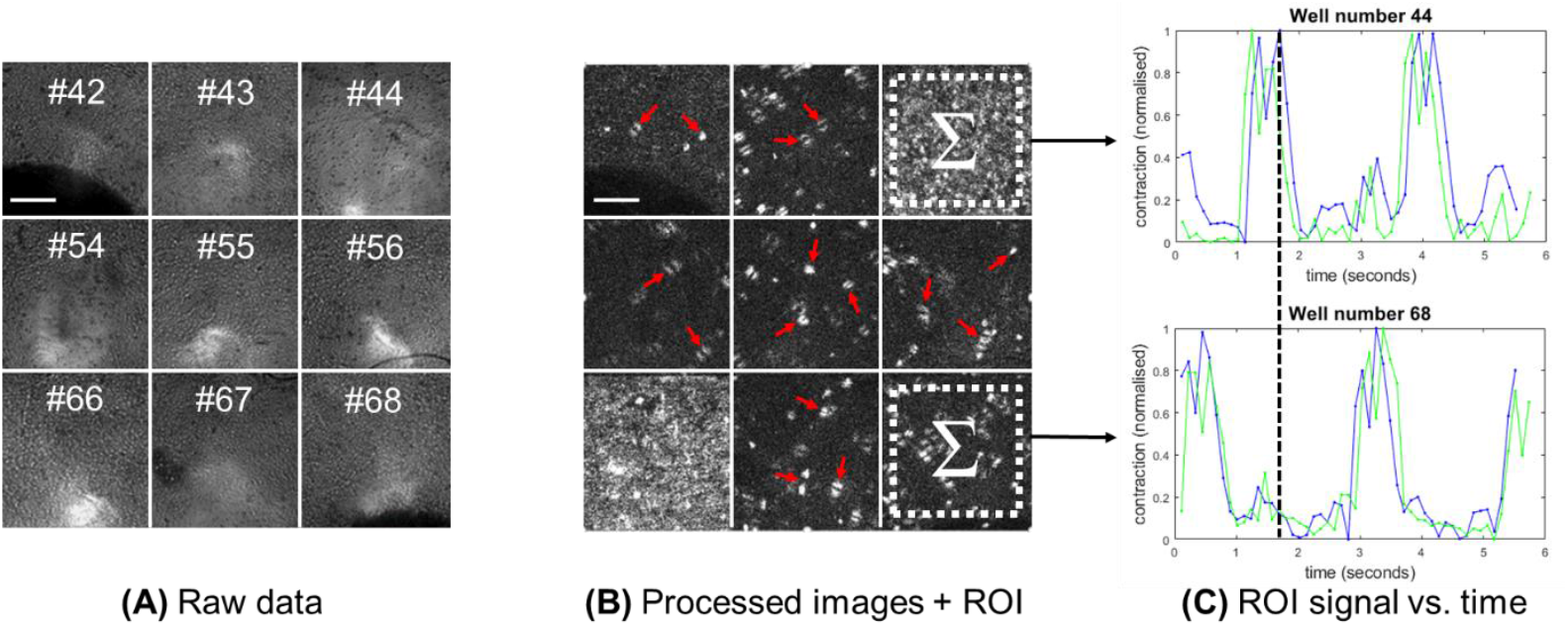
Demonstration of the PV image processing technique. (A) Raw data acquired from nine wells in a 96 well plate. (B) The processed data at the time point shown by the dashed vertical line in (C). This is at the point that wells #44 and #66 are undergoing contraction. Contraction speed traces, produced by summing the signal within the ROI of the processed frames, are shown in (C). This shows that well #44 is undergoing peak contraction and well #68 is quiescent. False signal from bubbles (some of which are indicated by arrows) increases the level of background noise. Scale bars 500 µm.

## RESULTS

### Clyde Biosciences data

The high magnification, high resolution CB data was analysed using both PV and MM algorithms, with contraction speed profiles for each of the nine wells considered shown in Figure 5 with a detailed view shown in Figure 6. All the wells were recorded for the same 7 second period, capturing between 5 and 7 contraction events. Each of the contraction speed profiles have been normalised to provide a clearer comparison. A frame delay of 20 frames (corresponding to approx. 20% of the total contraction duration) was used to calculate the traces. This frame delay value is much higher than the 2-5 frames recommended for 100 Hz data and was chosen to accentuate any differences between the PV and MM methods.

**Figure 5.**
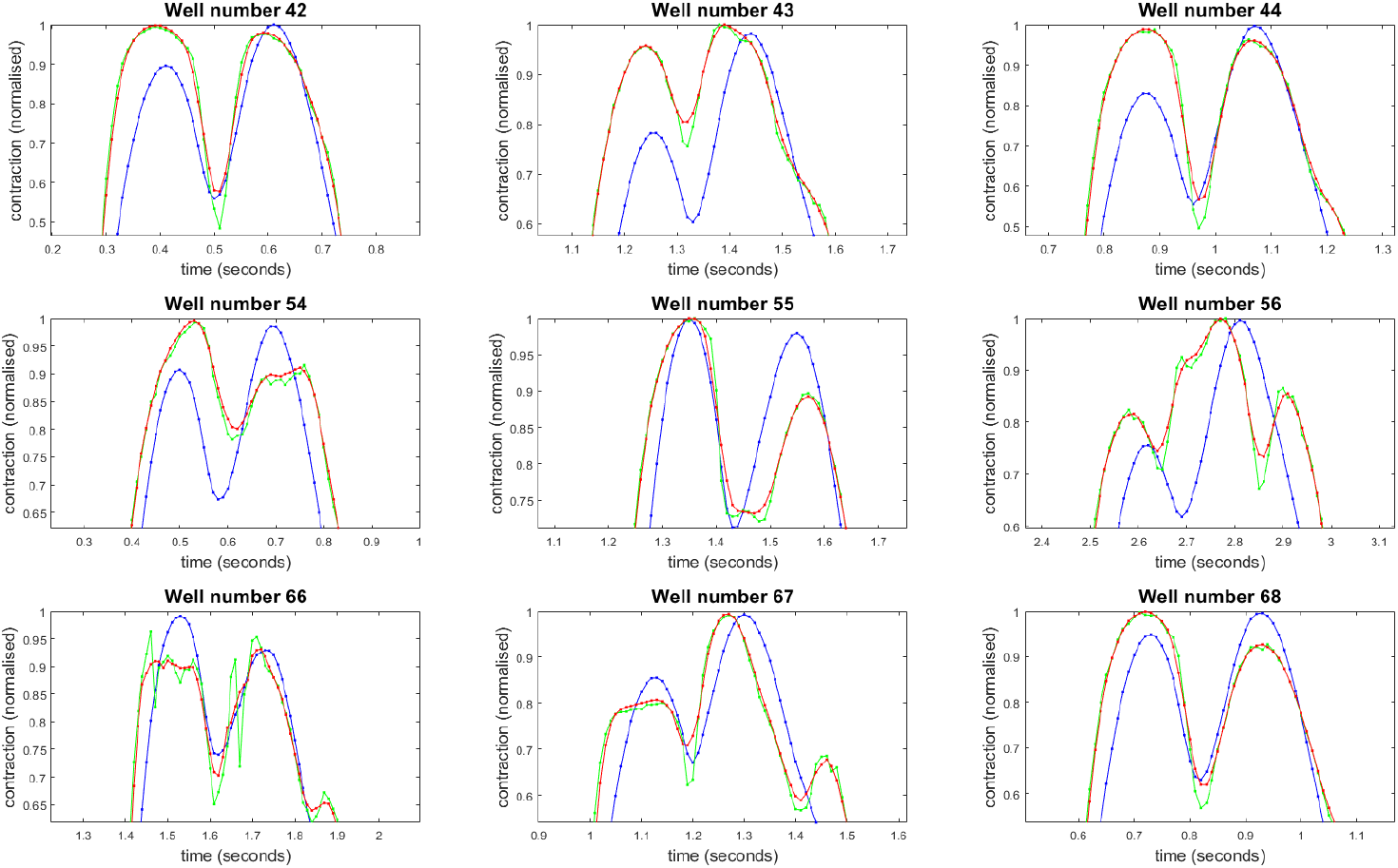
Detailed view of the first minima for the contraction speed curves shown in Figure 5, highlighting differences between the PV (blue) and MM (green) algorithms. For comparison, a four frame moving average of the MM data is also shown in red.

**Figure 6:**
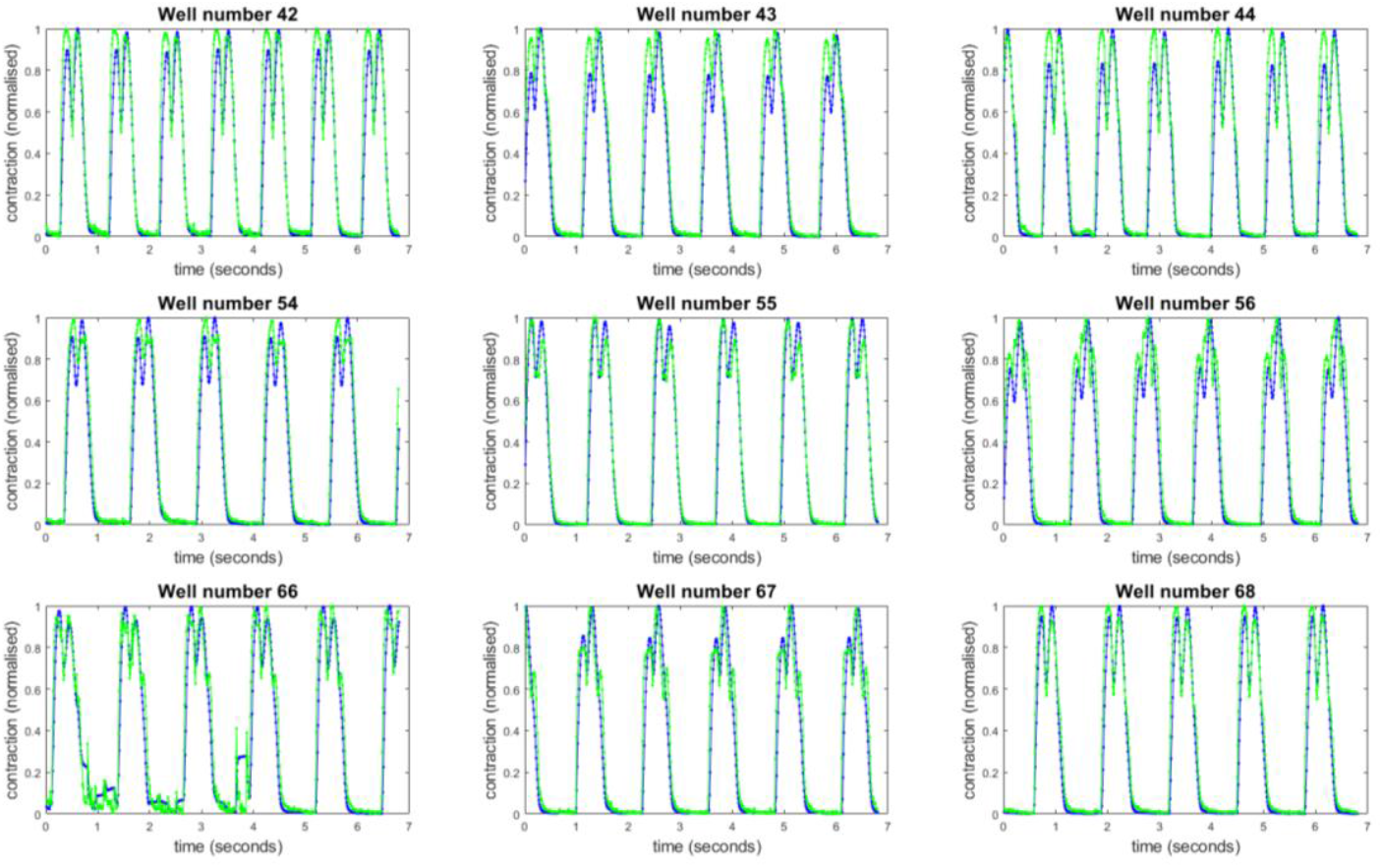
Comparison of the PV (blue) and MM (green) contraction speed data for image data collected on the Clyde Biosciences system. All curves have been normalised to the maximum peak value. Data acquisition took approximately 280 seconds

These traces were processed from spontaneous tissue contraction data in a controlled environment. The OKO labs stage incubator maintained a temperature of 37 degC, 95% humidity and 5% CO2. No drugs were added to the tissues before data collection and any differences observed in beat frequency or contraction speed between the nine wells under study are due to natural variations. The spontaneous beat rate can be seen to vary between 0.7 Hz and 1 Hz.

Figure 6 shows that the traces produced by the PV and MM algorithms closely follow each other. Qualitatively, the PV curves are smoother, with little difference in rise time and fall times. The minima that mark the points of greatest contraction align well for both PV and MM metrics, although in some cases there are triplet peaks in the contraction speed data for MM which are not seen with PV. These figures demonstrate that the variability data measured by the PV algorithm is a fundamentally different quantity to the absolute difference. To further indicate that the variance is not simply smoothing the MM data, a four-frame moving average of the MM data was calculated for comparison (red line, Figure 6). As the moving average is centred on the averaging window, no delay is introduced into the main features of the curve.

The PV and MM algorithms respond differently to the presence of vibration in the acquired image sequence. This is seen in the first half of the acquisition for well 66, where the PV metric registers a larger response than the MM. One reason for this is that the variance calculated from rapidly changing pixel values is unbounded, whereas the absolute difference will always be restricted to the 16-bit range of pixel values within the local area.

Both PV and MM algorithms calculate absolute contraction as well as contraction speed. This requires the use of a fixed reference frame against which to compare the current frame. The reference frame is calculated automatically by MM and will typically fall within the quiescent period of the monolayer contraction. For PV, there must be a comparison of pixel variance between the current and reference time points. This is achieved by calculating the pixel variance for a fixed number of frames adjacent to these time points and calculating the absolute difference of these values. Both PV and MM metrics will scale in response to temporal changes in the overall average image brightness. MM measures the average frame brightness and uses it to compensates the contraction amplitude data. This is harder to do for PV data as the scaling is not linear. For this reason the PV curves have not been corrected for baseline changes in image brightness.

### Nine well Multiscope data

Multiscope image acquisition of the nine wells was fully automated, sampling each well in an R, G, B colour sequence before moving on to the next well. Capturing images at 100 fps, the sampling rate per well and per colour was (100 / (9*3)) = 3.7 fps. Before the samples could be imaged, they were removed from the stage incubator and transferred to the Multiscope. As the cardiomyocyte monolayers cool down, the beat frequency reduced significantly from 1 Hz to 0.25 Hz.

The contraction speed profiles can be seen in Figure 7 for both the MM (green) and PV (blue) algorithms. These data were extracted from images acquired under red illumination, where the focus was sharpest (see Appendix A). Due to the lower sampling rate, a 2-frame delay was implemented, resulting in a much smaller distinction between the two processing algorithms. Despite the two orders of magnitude reduction in sampling frequency per well compared to the CB data, two or three contraction peaks can be clearly seen in all nine wells. In many cases, the dip in speed upon reaching maximum contraction can also be resolved within each contraction peak.

**Figure 7:**
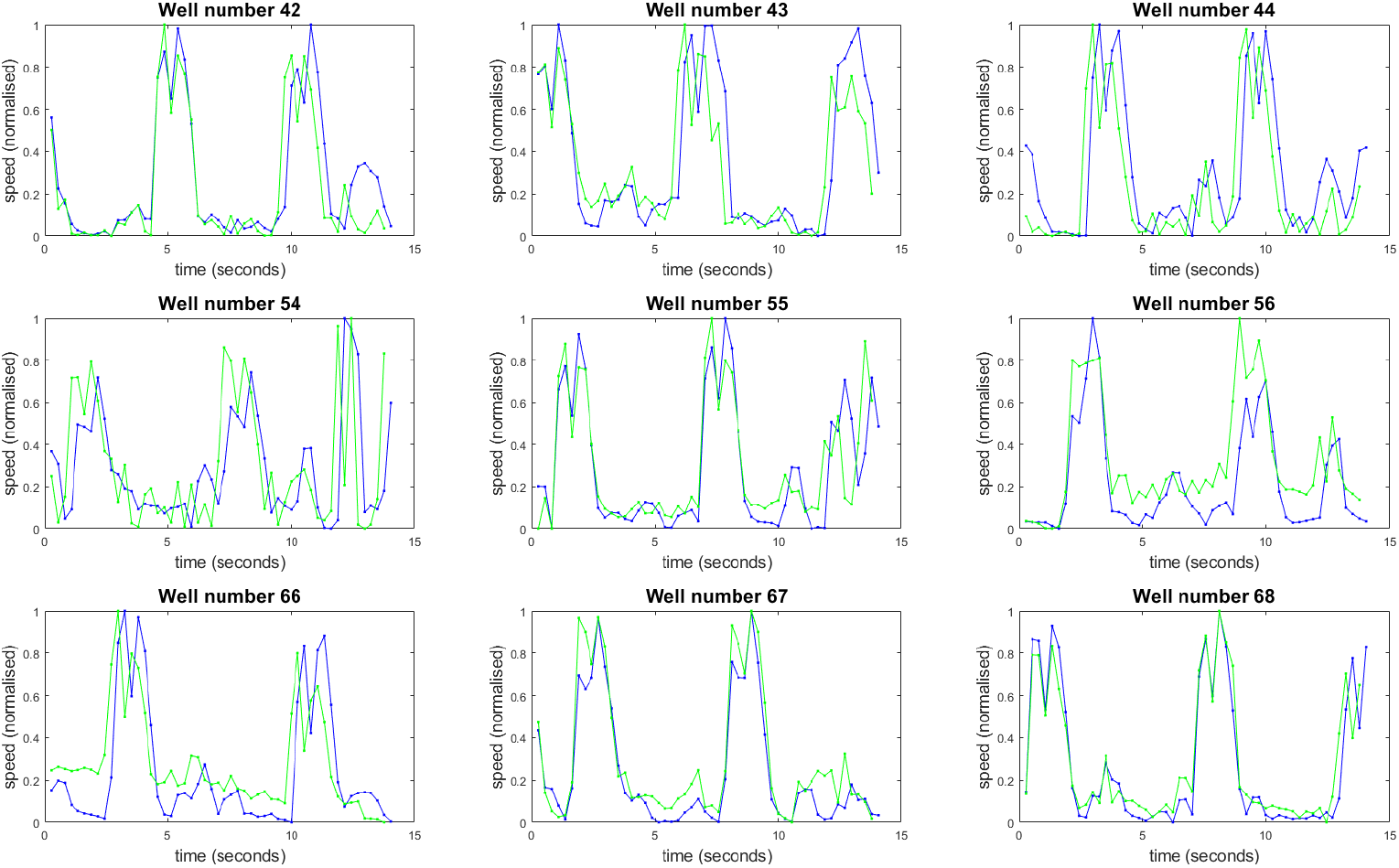
Contraction speed traces extracted from each of the nine wells imaged using the Multiscope. The traces were analysed using both PV (blue) and MM (green) algorithms. Data acquisition took a total of 15 seconds.

In all cases the data was acquired in untreated tissue as a baseline measure of the beat rate before introducing drugs that may affect the contraction properties of the monolayers. It is clear from this data that a much higher sampling rate would be required in order to provide the sensitivity to small, changes in the contraction profile that may result from an intervention.

To increase sampling rate per well, a two-stage approach might be suggested in which data are acquired in 3 colours for each well in stage 1 and the data used to determine the which colour is in best focus for each well. Once this has been determined, stage 2 acquisition can begin with only one LED colour per well and sampling each well sequentially at 100 Hz for several contractions before moving on to the next well. By emulating the CB protocol in this way, it would retain the benefits of high temporal resolution whilst gaining in the automation of the data acquisition. Cycling through each of the wells sequentially is expected to take ∼5 seconds per well, 45 seconds for nine wells, or 8 minutes for all 96 wells. Revisiting each well within 10 minutes would be more than sufficient when looking for longitudinal physiological responses to drug treatments over many hours or days.

### Single well Multiscope data

To compare the effect of sampling rate on the MS measurement of the contraction profile, an additional data set was acquired with a sampling rate of 50 Hz. This was performed for well #42 using a single colour channel (green). The results are shown in Figure 8.

**Figure 8:**
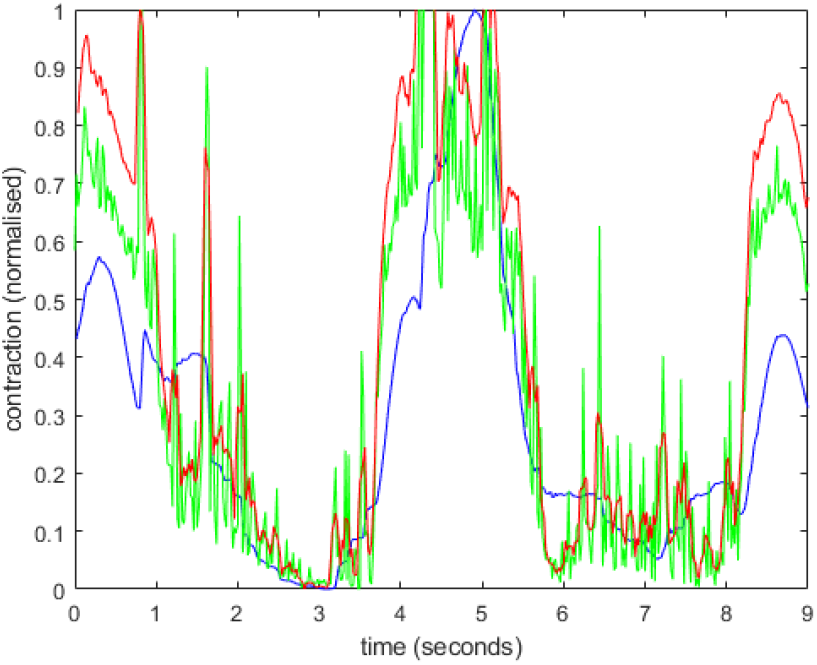
Contraction speed data from single well, single colour channel images captured by the MS at 50 Hz. Data processed using PV (blue), MM (green) and MM smoothed with a 4-frame moving average (red).

The data was taken with the MS on an undamped table which introduced vibrations to the sample. Vibrations appear as sharp spikes in the traces produced by the analysis software. As noted above, rapid cooling of the monolayer has greatly increased the contraction interval to 4 seconds. In response, the frame delay was doubled to N=40 frames. This value was applied to both PV and MS algorithms. For comparison, a smoothed version of the MM trace (using a four-frame box car average) is also shown in Figure 8. In this figure it can be seen that the PV trace is much smoother than the MM trace, although the rise time and fall times appear longer. Further data, over more contraction cycles, acquired on a vibration isolation table, would be required for a more rigorous analysis.

### Visualising tissue contraction

In both PV and MM algorithms, the processed images are summed to extract the contraction speed traces, losing all spatial information. For the PV method, it was possible to extract out the processed frames individually and identify their evolution in time. Two of these frames were extracted for a single contraction event, corresponding to the two peaks in the contraction speed trace. These images were then colour coded for each peak (red for first peak, blue for second peak) to identify the directions and amplitudes of the contraction displacement across the sample. The resulting image is shown in Figure 9. Contraction displacement is orthogonal to the ‘fringes’ visible in the image, with contraction amplitude indicated by the separation of the fringes. This image can provide useful information about the connectivity of the syncytium and how well bound the monolayer is to the well plate.

**Figure 9:**
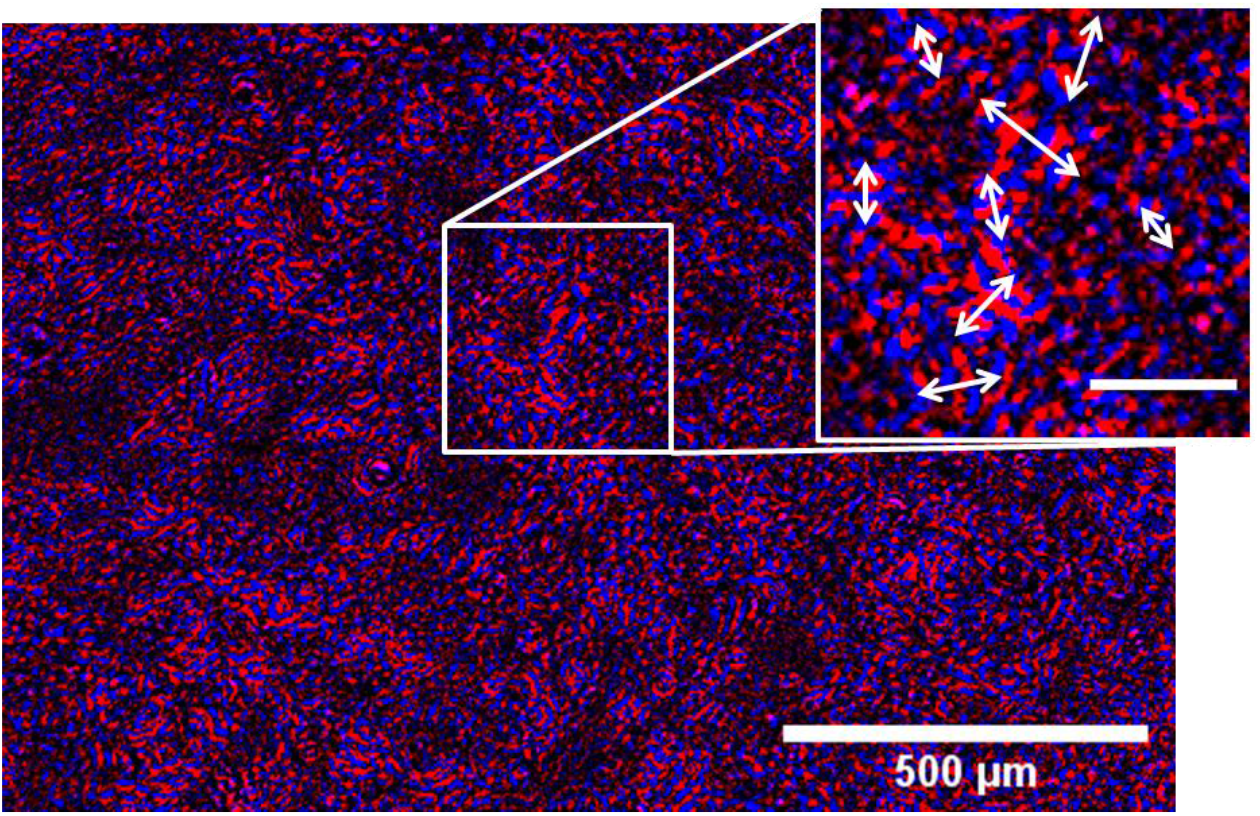
An overlap of two false colour PV processed images. The processed data is calculated at two different points in time (peak velocity of contraction, peak velocity of relaxation). The image shows the spatial pattern of contraction across the monolayer. The direction of motion is orthogonal to the red and blue fringes, with the amplitude of displacement given by the fringe spacing (larger spacing for larger amplitudes). Inset: arrows indicate the monolayer displacement direction and amplitude highlighting complex forces acting on the monolayer. Inset scale bar 100 µm

## DISCUSSION

This initial trial has demonstrated the potential of the unique Multiscope architecture to automate the acquisition and processing of contraction data in cardiomyocyte monolayers. Whilst the temporal resolution and acquisition duration in the current acquisition protocol is insufficient to accurately capture the contraction parameters commonly used for monolayer models, this can be addressed through changes in the acquisition protocol. Equally, averaging of the contraction profiles for a given well (routinely performed on CB data) was difficult to perform for this reason. Again, all these parameters can be easily changed through control software, which may begin with automated determination of the best colour to use in each well. Alternatively, acquiring at >100 Hz for > 5 seconds in a single colour for each well before switching colour could achieve the same result.

The lack of a stage incubator on the Multiscope was a major barrier to obtaining accurate contraction data. This could be achieved by making adaptations to the existing well plate holder design to incorporate the dimensions of the stage incubator. This would enable longitudinal imaging of the well plate before and after a drug intervention.

The PV algorithm was found to improve the smoothness of the traces extracted from both the CB and MS data without any noticeable compromise in responsiveness. This is shown in Figure 6, where the timing of the inflection points is largely similar for both algorithms. Reducing noise through the application of the PV algorithm is expected to increase sensitivity to introduced pharmacological changes in the contraction profiles.

The use of strongly collimated light greatly improved image contrast and sensitivity to tissue contraction. One negative effect of the collimation was introducing sensitivity to phase changes outside of the depth of focus. Meniscus effects, bubbles, circulating scatterers all acted to increase the level of background variability which masked the tissue contraction. An inverted Multiscope design would allow for objective lenses with focal lengths shorter than the well depth, increasing the NA and reducing the depth of field which would reduce the influence of out-of-plane artefacts. At higher magnifications, contrast from cell boundaries would also become clearer, potentially improving further sensitivity to tissue movement, albeit at the cost of very localised sampling.

Finally, in this experiment, data was acquired on a standard office desk, located in the same room as the CB microscope. This introduced background vibrations and the occasional jolt, the latter of which could produce trace peaks as large as the tissue contraction itself. Whilst jolts in the data are easy to identify and mitigate as they appear in all wells the same time, the use of a floating optical table would be beneficial to stop them arising in the first place.

## CONCLUSION

We have implemented a new type of parallelised microscope for the acquisition of contraction data from cardiomyocyte monolayers. We have demonstrated that the ‘Multiscope’ can acquire contraction data from nine wells in a 96 well plate by illuminating them in rapid succession without the need for any moving parts. To collect seven seconds worth of contraction data for nine wells took seven seconds. This can be compared with 280 seconds for the motorised stage, due to the need to first manually focus and align the centre of the field of view on each well, in addition to the time required for motorised stage translation between wells. Whilst the temporal sampling per well is lower on the Multiscope, this represents an increase in speed by a factor of 40.

The use of an adapted image processing algorithm (‘Pixel variance’) has shown that it is capable of significantly reducing noise in the extracted contraction traces without compromising the key timing points. In future work we will look to extend the Exeter Multiscope prototype further to enable automated contraction measurements across all 96 wells at 100 Hz acquisition rates per well.

## APPENDIX A

### Optimising Multiscope analysis parameters

Before processing the MS image data, three parameters first needed to be determined: the frame delay, window size and focal depth. The frame delay determines the time interval over which the raw image data are processed (Figure 3). For the CB data, this was set at around 20% of the interval between contraction. For the MS data, there are approximately 10 sampling points between peaks, meaning the frame delay is expected to be around 2 frames. The optimal frame delay is that which maximises the signal to background ratio of the peaks in the contraction speed trace. The signal to background ratio was calculated as the ratio of the highest 10% of values in the trace to the lowest 10% of values in the trace. The same data set, composed of all nine wells, was processed with a 1-, 2-, 3-, 5-, and 10-frame delay. The average signal to background ratio was calculated for all frame delays. The results are shown in Figure 10(A), which indicates that there is a marginal advantage to using a 2-or 3-frame delay. In this case it was decided to use a 2-frame delay (= 0.54 s) for all MS data calculations.

**Figure 10:**
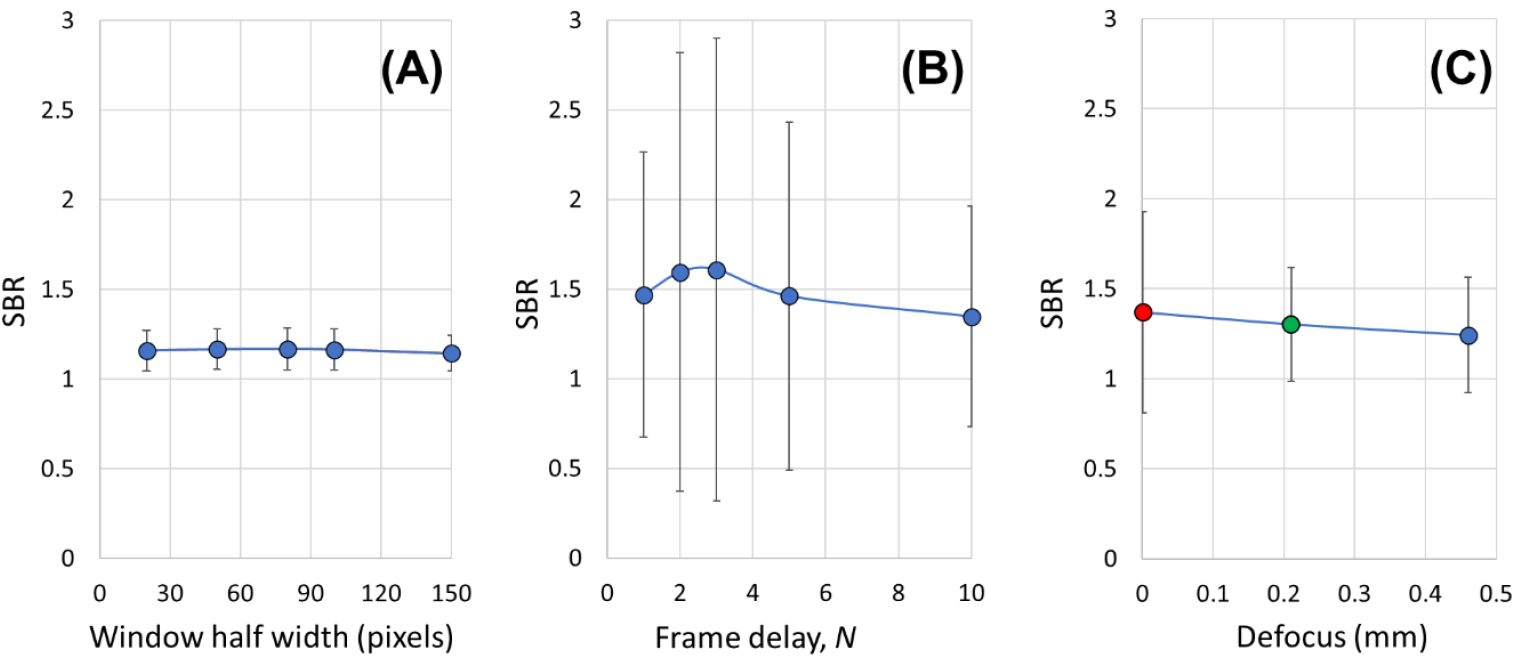
Variation in contraction signal contrast as a function of cropping window size (A), frame delay and (C) illumination wavelength.

Second, the illumination wavelength that produced the best sample focus was determined by using the SBR as before, but this time comparing the three colour channels for all nine wells. The results indicated that the focal plane of the red channel was closest to the monolayer sample, which corresponds well with a visual inspection of the data from each channel.

Finally, the width of the region of interest used to crop the raw image data from each well was determined. This was calculated for window half-widths of 25, 50, 80, 100 and 150 pixels. The main trade off here was ensuring that the window was large enough to maximise the signal contained within the processed frame, whilst being small enough to avoid any effects from the edges of the well. In this analysis, the first well (well #44) had to be removed from this analysis due to the presence of a bubble which interfered with the average calculation.

## APPENDIX B

### Mathematical comparison of PV and MM algorithms

This section compares the MM and PV algorithms in terms of the noise propagation. This is important to understand how differences in these output metrics originate and how they scale with the size of the frame delay, *N*. In this analysis we consider a single pixel value, *x*_*i*_, corresponding to frame *i* (see Figure 3). For brightfield imaging, we assume that the photon statistics are dominated by shot noise (Poisson distribution), whereby the pixel values recorded on the camera are taken from a distribution with mean and variance values equal to *x*_*i*_.

Both MM and PV algorithms quantify the change in pixel value over a pre-defined frame delay (*N* frames). The MM metric calculates the modulus of the difference, whereas the PV calculates the standard deviation of the pixel values over the interval. We want to identify how the errors in *x*_*i*_ propagate to the metric used in each case.

Beginning with the MM metric, we define this as:

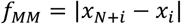

For which the error is propagated as:

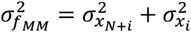

Which, if the pixel values are close to each other (*x*_*N*+*i*_ ≈ *x*_*i*_) we can simplify to:

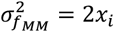

For the PV metric, we begin by calculating the variance (*f*_*VAR*_) of the pixel values over time:

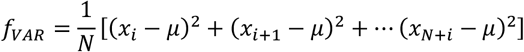

Where *μ* is the mean pixel value over the range *x*_*i*_ to *x*_*N*+*i*_. We employ the error propagation formula:

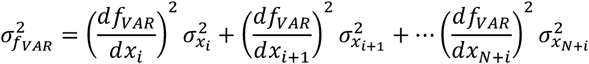

Where the partial derivative in each of these terms has the form: 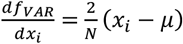

Making the same assumption that the pixel values are not very different between frames, we can write:

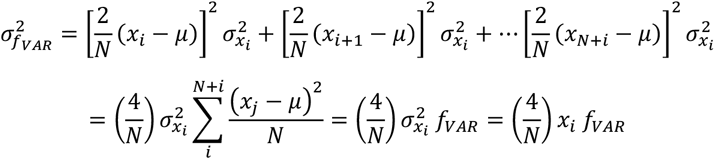

Finally, we can relate the error in the pixel variance, 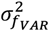, to the error in the standard deviation of the pixel values,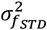. Given that 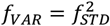, we can write:

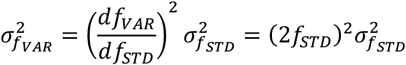

Making *f*_*STD*_ the focus of this equation we have:

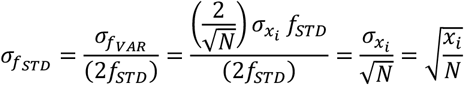

We can then write down the ratio of the standard error for these two metrics as:

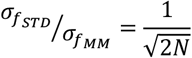

This predicts a 2X reduction in noise in the PV metric for the MS data (N=2) and a 6.3X reduction in noise for the CB data (N=20) relative to the MM metric. Whilst the improvements are harder to see in the MS data, there are clear differences in the CB data between the MM (green) and PV (blue) traces (Figure 8).

## ACKNOWLEDGEMENTS

The authors would like to acknowledge the generous contribution of time, people, tissues and reagents by Clyde Biosciences that made these measurements possible.

## AUTHOR CONTIBUTIONS

Multiscope design and construction (DWH, ADC), experimental design (ADC, GS, GB, CM), data collection (ADC, TW, SG, LH), data analysis (ADC, FB), mathematical modelling (ADC), ZEMAX simulation (SM), first manuscript draft (ADC), manuscript revisions (CM, SM, GS, ADC), funding acquisition (SM, CM, FB, GS).

